# HIF1α controls adipose tissue growth through metabolic rechanneling

**DOI:** 10.1101/2024.05.28.596336

**Authors:** Chaonan Zhu, Meiqian Wu, Minh Duc Pham, Yue Wang, Arka Provo Das, Yijie Mao, Peter Mirtschink, Ting Yuan, Jaya Krishnan

## Abstract

Excessive expansion of white adipose tissue occurs when energy intake exceeds demand. It creates a state of relative hypoxia that is directly linked to activation of the hypoxia-inducible factor (HIF) in obese human subjects and mouse models of obesity. Whether HIF1α function is important for the development of basal adiposity and diet-induced obesity remains unclear. In the present study, we genetically ablated Hif1α in adipocytes and analyzed the corresponding mice for both basal and diet-induced obesity-associated visceral white adipose tissue mass as well as parameters of systemic glucose homeostasis and peripheral insulin sensitivity. We found that inactivation of Hif1α in mouse adipocytes inhibited basal adiposity and suppressed nutrient-overload-induced adipose tissue growth. These changes in adiposity were associated with improved systemic glucose homeostasis and peripheral insulin sensitivity. Mechanistically, Hif1α mediated effects on adipocyte metabolism by re-routing glycolytic intermediates into the glycerolipid shunt, leading to increased de novo triacylglyceride synthesis in hypertrophic adipocytes. Our results have established key roles for HIif1α in controlling adipocyte growth and metabolism and adipose tissue expansion under basal conditions and in response to a high-fat diet, highlighting the central role of hypoxia and HIF1α in the development of obesity.

**Article Highlights:**

- Hif1α function is critical for basal adipose tissue mass expansion, and is also required for fat accumulation in response to high fat diet.
- Co-option and co-activation of the glycerolipid biosynthetic pathway by glycolysis is an important modulator of visceral adiposity.
- These findings reveal an important role of Hif1α in the adipose tissue through the coupling of glycolysis and lipid anabolism in response to HFD, which may provide a potential therapeutic target for type 2 diabetes in the future.

## Introduction

Obesity is a complex metabolic disorder linked to an increased risk of developing common human diseases including type 2 diabetes, hypertension, dyslipidemia, cardiovascular disease and cancer, and is a consequence of excessive expansion of white adipose tissue (WAT) (1–3). More specifically, it is the increase in visceral fat mass that is the best predictor of obesity-related morbidity (1). WAT expansion in obesity could be mediated by hypertrophy (enlarged adipocytes) and/or hyperplasia (increased number of adipocytes) (4). Although the increases in both adipocyte cell size and cell number contribute to adipose tissue growth, the former is considered to be strongly influenced by diet. Indeed, nutrient overload, a key factor in promoting the obese state, induces adipocyte hypertrophy and can promote the development of insulin resistance and glucose intolerance (5–8).

Visceral adipose tissue (VAT) of obese humans and mouse models of obesity is commonly poorly oxygenated (2), which is at least in part due to an inability of the pre-existing adipose tissue vasculature to meet the oxygen demands of the expanding adipose mass resulting from chronic and excessive nutrient consumption (9). Decreased tissue oxygenation (hypoxia) is, in mammalian cells, invariably associated with the activation of HIF, dimers composed of oxygen-regulated HIF1α, HIF2α or HIF3α subunits and a constitutively expressed HIF1β (also known as ARNT) subunit, and activation of HIF target genes. The products of HIF target genes mediate the induction of adaptive cellular processes, including the promotion of angiogenesis, regulation of cell proliferation and survival, as well as alterations in cellular metabolism (10). Evidence suggests that among the HIFα subunits, HIF1α preferentially activates genes important for glycolysis, whereas HIF2α appears to favor the activation of genes involved in angiogenesis (11).

Several lines of evidence suggest that HIF1α accumulates in adipocytes of obese humans and mouse models of obesity (12–17), and that increased HIF1α protein leads to increased adipose fibrosis in obesity (18,19). Functional studies of HIF1α in mouse WAT either through means of overexpression of a constitutively active HIF1α (12), expression as a dominant-negative form (18,20), the deletion of the gene in normotrophic white adipose tissue (14) or its inducible ablation in hypertrophic adipocytes (15), have revealed a central role for this transcription factor in promoting fat accumulation and associated pathologies including glucose intolerance and insulin resistance. However, the mechanisms through which HIF1α promotes the development of excessive fat accumulation are only beginning to be understood.

In this regard, it has been reported that the development of cardiac hypertrophy is associated with a HIF1α-dependent reprogramming of cellular metabolism that encompasses a switch from fatty acid utilization to glycolysis and increased synthesis of free fatty acids (FFA) and triacylglycerides (TAG) (21,22). In this context, the switch to a glycolytic form of metabolism can be attributed to the activation of genes whose products encode glucose transporters, glycolytic enzymes, lactate dehydrogenase A (Ldh-A) and pyruvate dehydrogenase kinase 1 (Pdk1), which are involved in shunting pyruvate away from mitochondria (23). On the other hand, HIF1α directly activates the transcription factor peroxisome proliferator activated receptor γ (PPARγ), which is known to promote lipid anabolic processes. Thus, by directly activating glycolytic and indirectly lipid anabolic pathways, HIF1α promotes the lipid accumulation in cardiomyocytes in response to pathological stressors known to induce cardiac hypertrophy (21,22,24). Given that HIF1α supports efficient glucose to lipid conversion in the context of cardiac hypertrophy, we asked whether this capacity of HIF1α is also a critical component of white adipocyte growth and metabolism.

Altered or dysfunctional metabolite channeling has been shown to play a regulatory role in DNA replication, cell proliferation and survival. However, it is unclear whether rechanneling of metabolites facilitates transition to obesity and how these rechanneling events are regulated in expanding adipocytes. Therefore, we embarked on defining how flux along key metabolic pathways are differentially regulated between normal and hypertrophic adipocytes and if dysregulated metabolic flux imposes a regulatory role in basal adiposity. Hence leading to dysregulated metabolite buildup and subsequent rechanneling the role of Hif1α in basal– and dietary obesity-associated visceral white adipose tissue mass expansion and its effects on adipocyte metabolism as they relate to rechanneling of glycolytic intermediates into the glycerolipid shunt and de novo triacylglyceride synthesis in hypertrophic adipocytes.

## Methods

### Animal Breeding and Maintenance

Hif1α^fl/fl^ mice were obtained from Randall S. Johnson (University of California, San Diego, USA). The Fabp4/aP2-cre/+ line was generously provided by Ronald M. Evans (University of California, San Diego, USA) (25). The data presented in this manuscript represents studies with male mice, maintained at the RCHCI, ETH-Zurich specific pathogen-free facility. Plasma glucose levels were monitored at the beginning of the night phase for the random-fed group and the consecutive morning after a 12-h food deprivation for the fasted group. Maintenance and animal experimentation were in accordance with the Swiss Federal Veterinary Office (BVET) guidelines.

### GTT and ITT Measurements

Mice were fasted overnight, and glucose levels were determined 30 min before glucose injection. After an intraperitoneal injection of 1.0g of glucose/kg bodyweight, glucose levels were determined with an AccuCheck Aviva glucometer (Roche) at 0, 15, 30, 60, 90, 120 and 150 min using blood from the tail vain. For the ITT, after an intraperitoneal injection of 0.75U of insulin/kg bodyweight (Actrapid, Novo Nordisk), glucose levels were determined at 0, 15, 30, 60 and 90 min as described above. Mice were statistically analyzed with Excel (Microsoft) and areas under curve (AUC) were calculated. All GTT and ITT measurements were performed on 10-15 mice of the respective genotype and diet, unless indicated otherwise.

### Immunofluorescence of Cryosections

Tissues were embedded in OCT (Sakura Finetek). Cryosections were prepared as described (26) and fixed for 10 minutes with 4% paraformaldehyde (PFA), permeabilised by incubation with 0.2% Triton X-100 for 10min and immunofluorescence reactions and mounting of specimens was performed as described previously (26). Oil Red O staining was performed and quantified as previously described (22), while BodiPy staining was done with the HCS LipidTox Neutral Lipid Stain (Invitrogen), as recommended by the manufacturer.

### Cell Size and Cell Number Quantification

Micrographs were taken on wide-field microscopes (Leica) with a color CCD camera. Adipocyte size was quantified with the ImageJ software on three independent WAT sections per animal and 30-60 adipocytes were quantified per section. Adipocyte number was determined by counting primary adipocytes isolated from visceral fat pads of mice of the respective genotypes.

### Antibodies and Fluorescent Reagents

Hif1α antibodies were obtained from Santa Cruz Biotechnology. Phalloidin, BodiPy and DAPI were used as recommended by the manufacturer (Invitrogen).

### Immunoblotting and Immunoprecipitations

Dissected adipose tissue was homogenized by freeze slamming and solubilized in a modified SDS sample buffer sonicated and boiled for 5min. Protein extracts were resolved on 10 or 15% polyacrylamide minigels (BioRad) and transferred overnight onto nitrocellulose membrane (Amersham). Immunodetection and visualization of signals by chemiluminescence was carried out as described (26). Immunoprecipitation for Pgc1α was performed as previously described (27) Densitometric analysis of immunoblots were performed with ImageJ.

### Quantitative RT-PCR

Total RNA was isolated with Trizol (Invitrogen) and cDNA was generated using Superscript II (Invitrogen) from individual hearts as recommended by the manufacturer. qPCR reactions were setup as recommended by the manufacturer (Roche) and analyzed on the Roche LightCycler 480.

### Glycolytic Flux Measurements

Cellular glycolytic flux was measured using a Seahorse bioscience XF24 analyzer in 24 well plates at 37°C, with correction for positional temperature variations adjusted from four empty wells evenly distributed within the plate. Primary adipocytes were seeded at 20000 cells per well and washed and 630 μl of non-buffered (sodium carbonate free) pH7.4 DMEM supplemented with 10% fetal calf serum was added to each well, prior to measurements. After a 15 minute equilibration period, three successive 2 minute measurements were performed at 5 minute intervals with inter-measurement mixing to homogenize oxygen concentrations in the medium and each condition was measured in five replicates in independent wells. Glycolytic flux was measured as described by the manufacturer. Briefly, freshly glucose was injected in glucose-free minimal medium and glycolytic rate was calculated as the increase extracellular acidification (ECAR) above baseline.

### Enzyme Activity and Substrate Content Measurements

Spectrophotometric measurements were performed using a Spectra MAX 190 plate reader (Molecular Devices). After reaching steady state conditions, the starting reagents were added and absorption changes were registered every 10 s over a period of 20 to 30 min. Activities were calculated from the linear phase. For glyceraldehyde-3-phosphate dehydrogenase activity measurement, lysates were assayed in glycine (100 mM, pH 8.5), 1 mM DTT, 0.5 mM EDTA, 2.2 mM glyceraldehyde-3-phosphate and NAD+. The reaction was started with the addition of 10mM arsenate and formation of NADH was monitored at 340 nm at 30°C. Glycerol-3-phosphate dehydrogenase activity measurements were performed with lysates that were assayed in glycine (100 mM, pH 8.5), 0.5 mM EDTA, 1 mM DTT, and 1 mM dihydroxyacetone-phosphate. The reaction was started with the addition of 1 mM NADH and the formation of NAD+ was monitored at 340 nm at 30°C. Mitochondrial activity was assessed as described (28). Lysates were assayed in glycine (100 mM, pH 8.5), 0.5 mM EDTA, 1 mM DTT, 0.1 mM palmitoyl-CoA, and 0.1 mM 5,5’-dithio-bis(2-nitrobenzoic acid) for glycerol-phosphate acyltransferase activity measurements. The reaction was started with the addition of NADH and the formed DTNB-CoA complex was measured spectrophotometrically at 412nm. Glycerol-3-phosphate content was performed as described previously (29). Dihydroxyacetone phosphate levels were measured by incubating lysates in 100 mM glycine, pH 8.5 in the presence of 2.3U glycerol-3-phosphate dehydrogenase. The reaction was started by the addition of 1mM NADH. Absorbance changes were monitored at 340 nm.

### Statistical Analysis

All statistical analyses and tests were carried out using Excel software (Microsoft). Data are represented as mean and error bars indicate the standard error of the mean (±SEM). Significance was assessed by unpaired two-tailed t-test.

## RESULTS

### Obesity drives hypoxia and Hif1α activation in visceral adipose tissue

Nutrient overload is linked to accelerated adipose mass expansion and the development of intra-adipose hypoxia (2). To confirm the relationship between nutrient overload and accelerated adipose mass expansion, with the accumulation of HIF1α in adipocytes, wildtype C57BL/6J mice were maintained on a high-fat diet (HFD). HFD maintained mice revealed increased incorporation of the hypoxia marker pimonidazole and augmented Hif1α protein accumulation (Figures 1A and 1B), concomitant with increased body weight, adipose cell size and increased triacylglyceride (TAG) levels (Figures 1C-1F), compared to adipose tissue of mice fed a normal chow diet (NCD). These data imply that adipose mass expansion in response to a HFD is characterized by adipose tissue hypoxia and the accumulation of Hif1α in adipocytes, consistent with earlier results (13,15).

**Figure 1.**
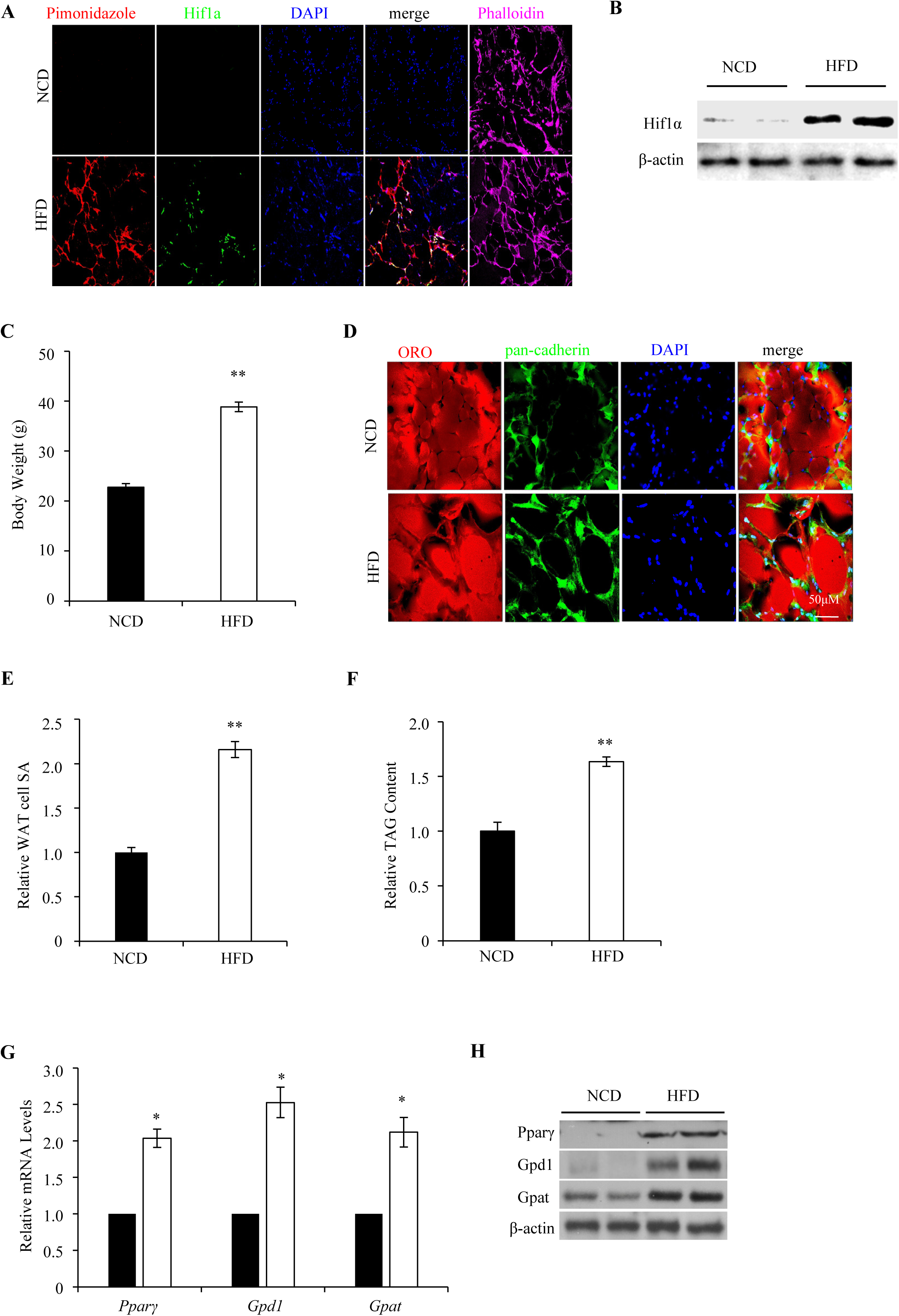
Obesity drives hypoxia and Hif1α activation in visceral white adipose tissue. (A) Pimonidazole was injected into mice maintained on either a NCD or HFD. Visceral white adipose tissue (WAT) sections of these mice were assessed for pimonidazole incorporation (red) by indirect immunofluorescence microscopy. Sections were counterstained with anti-Hif1α antibody (green), DAPI (blue) and phalloidin to mark actin (pink). (B) Western blot shows the Hif1α protein expression levels in isolated WAT from NCD– and HFD-fed C57BL/6J mice. (C) Body weight is expressed as means ± SEM from NCD– and HFD-fed mice (n=6), Data are expressed as mean ± SEM; \*\**P* < 0.01 vs. NCD, all by unpaired two-tailed Student’s t-tests. (D) WAT sections of NCD– and HFD-fed mice were stained with the neutral lipid marker, oil red O (ORO, red), and DAPI (blue), and counterstained with the cell boundary marker, pan-cadherin (green), and analyzed by immunofluorescence microscopy. Scale Bar: 50 μM. (E) Quantification of visceral WAT cell surface area (SA) of NCD-and HFD-fed mice. Data are expressed as mean ± SEM (n=7). \*\**P* < 0.01 vs. NCD, all by unpaired two-tailed Student’s t-tests. (F) Quantification of visceral WAT TAG content of NCD-and HFD-fed mice. TAG content was normalized to tissue weight. Data are expressed as mean ± SEM (n=5). \*\**P* < 0.01 vs. NCD, all by unpaired two-tailed Student’s t-tests. (G) Relative mRNA expression levels of the *Pparγ*, *Gpd1* and *Gpat* in isolated WAT from NCD– and HFD-fed C57BL/6J mice. Data are expressed as mean ± SEM (n=3). \*\**P* < 0.01 vs. NCD, all by unpaired two-tailed Student’s t-tests. (H) Western blot shows the Pparγ, Gpd1 and Gpat protein expression levels in isolated WAT from NCD– and HFD-fed C57BL/6J mice. NCD, normal chow diet; HFD, high-fat diet; WAT, white adipose tissue.

Adipocyte TAG levels are determinants of adipocyte size and insulin sensitivity. Pparγ is a potent inducer of TAG synthesis and has been shown to be a downstream target of Hif1α in diseased cardiomyocytes (22). Due to the observed increase in the adipocyte size and TAG content of wildtype C57BL/6J maintained on HFD, we asked if Pparγ expression and transcriptional function was altered in white adipose tissue (WAT) of C57/Black6 mice. To that end, WAT biopsies of wildtype C57/Black6 mice subjected to the NCD and HFD protocol were assessed for expression of Pparγ, and its target genes, glycerol phosphate dehydrogenase 1 (Gpd1) and glycerol phosphate acyltransferase (Gpat). Gpd1 and Gpat are rate-limiting components of the glycerolipid biosynthetic shunt, and are important regulatory components of de novo TAG synthesis (30,31). As shown in Figure 1G and 1H, mRNA and protein levels of the respective genes were increased in WAT biopsies of mice maintained on HFD, compared to controls.

### Adipocyte specific Hif1α ablation protected from HFD induced obesity in mouse *in vivo*

While these findings establish a correlation between Hif1α accumulation, Pparγ expression and its downstream target genes Gpd1 and Gpat, it is unclear if Hif1α is functionally linked with the induction of Pparγ and its targets *in vivo*, and if Hif1α function is necessary for de novo glycerolipid biosynthesis in adipocytes as previously observed in diseased cardiomyocytes (22). To address this, we generated mice deficient for Hif1α specifically in adipocytes. Mice carrying loxP-flanked Hif1α alleles (Hif1α^fl/fl^) (22) were crossed to Fatty acid-binding protein 4 (Fabp4/aP2)-Cre recombinase (Cre) transgenic mice (25) to generate aP2-Cre; Hif1α^fl/fl^ (thereafter referred to as Hif1α cKO). The Fabp4/aP2-Cre transgene induces Cre-mediated recombination in white and brown adipocytes. Deletion of Hif1α was confirmed by PCR-mediated detection of the recombined alleles and led to a ∼90% downregulation of Hif1α mRNAs in white adipose tissue (WAT) as assayed by quantitative PCR (qPCR) (Fig. 2A). Further, Hif1α-target genes including Solute Carrier Family 2 Facilitated Glucose Transporter 1 (SLC2A1/Glut1), Aldolase A1 and Vascular Endothelial Growth Factor (Vegf) were similarly repressed, whilst the non-Hif1α target Solute Carrier Family 2 (Facilitated Glucose Transporter), Member 4 (SLC2A4/Glut4), was largely unaffected (Fig. 2A). Hif1α cKO mice were born at the expected Mendelian ratio, had comparable body weight to control Hif1α^fl/fl^ mice and did not exhibit any overt phenotype (Fig. 2B). Although levels of food consumption and fasting and post-prandial blood glucose levels were similar between Hif1α^fl/fl^ and Hif1α cKO mice (Fig. 2C,D), glycated-hemoglobin (HbA1c) levels, an average measure of blood glucose levels over a prolonged period of time, was consistently lower in Hif1α cKO mice compared to Hif1α^fl/fl^ control (Fig. 2E).

**Figure 2.**
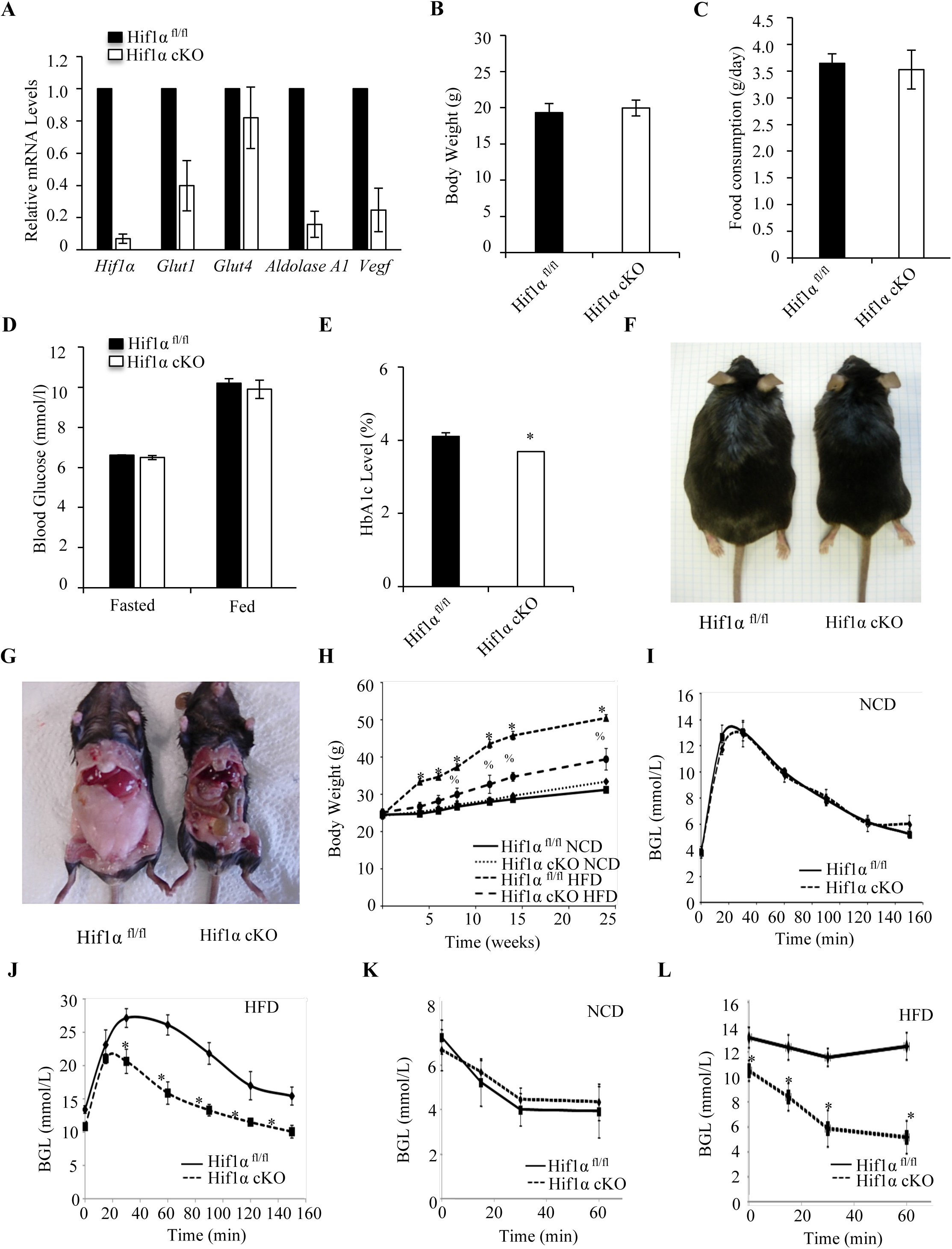
Adipocyte specific Hif1α ablation protected from HFD induced obesity in mouse *in vivo*. (A) Relative mRNA expression levels of the *Hif1α*, *Glut1*, *Glut4*, *Aldolase A1* and *Vegf* in HFD-fed Hif1α^fl/fl^ and Hif1α cKO mice. All values were normalized internally to 18S RNA expression and to the Hif1α^fl/fl^ control, respectively. Data are expressed as mean ± SEM (N=3). \**P* < 0.05 vs. Hif1α^fl/fl^, all by unpaired two-tailed Student’s t-tests. (B) Body weight for the Hif1α^fl/fl^ and Hif1α cKO mice maintained on HFD. Data are expressed as mean ± SEM (n=7). \**P* < 0.05 vs. Hif1α^fl/fl^, all by unpaired two-tailed Student’s t-tests. (C) Food intake for the Hif1α^fl/fl^ and Hif1α cKO mice maintained on HFD. Data are expressed as mean ± SEM (Hif1α^fl/fl^ n=5; Hif1α cKO n=5). all by unpaired two-tailed Student’s t-tests. (D) Fasting blood glucose and random fed blood glucose measurements in Hif1α^fl/fl^ and Hif1α cKO mice maintained on the HFD protocol. Data are expressed as mean ± SEM (Hif1α^fl/fl^ n=5, Hif1α cKO n=5.) all by unpaired two-tailed Student’s t-tests. (E) Glycated-heamoglobin (HbA1c) levels in the blood were measured n Hif1α^fl/fl^ and Hif1α cKO mice maintained on the HFD protocol. Data are expressed as mean ± SEM (Hif1α^fl/fl^ =5; Hif1α cKO n=8.) \**P* < 0.05 vs. Hif1α^fl/fl^, all by unpaired two-tailed Student’s t-tests. (F) and (G) Representative images of the Hif1αfl/fl and Hif1α cKO mice maintained on the HFD protocol. (H-L) Hif1α^fl/fl^ and Hif1α cKO mice were fed a NCD or HFD for 24 weeks. (H) Body weight is expressed as means ± SEM from Hif1α^fl/fl^ NCD n= 5; Hif1α cKO NCD n= 5; Hif1α^fl/fl^ HFD n= 7; Hif1α cKO HFD n= 7; \**P* < 0.05 vs. Hif1α^fl/fl^ NCD, %*P* < 0.05 vs. Hif1α^fl/fl^ HFD, all by unpaired two-tailed Student’s t-tests. (I) Glucose tolerance test (GTT) was measured in Hif1α^fl/fl^ and Hif1α cKO mice maintained on a NCD protocol. Data are expressed as mean ± SEM (Hif1α^fl/fl^ NCD n= 5; Hif1α cKO NCD n= 5). all by unpaired two-tailed Student’s t-tests. (J) Glucose tolerance test (GTT) was measured in Hif1αfl/fl and Hif1α cKO mice maintained on the HFD protocol. Data are expressed as mean ± SEM (Hif1α^fl/fl^ HFD n= 7; Hif1α cKO HFD n= 7). \**P* < 0.05 vs. Hif1α^fl/fl^ HFD, all by unpaired two-tailed Student’s t-tests. (K) Insulin tolerance test (ITT) was measured in Hif1αfl/fl and Hif1α cKO mice maintained on the NCD protocol. Data are expressed as mean ± SEM (Hif1α^fl/fl^ NCD n=5; Hif1α cKO NCD n=5). all by unpaired two-tailed Student’s t-tests. (L) Insulin tolerance test (ITT) in Hif1αfl/fl and Hif1α cKO mice maintained on the HFD protocol. Data are expressed as mean ± SEM (Hif1α^fl/fl^ HFD n=7; Hif1α cKO HFD n=7). \**P*< 0.05 vs. Hif1α^fl/fl^ HFD, all by unpaired two-tailed Student’s t-tests. NCD, normal chow diet; HFD, high-fat diet; WAT, white adipose tissue; BAT, brown adipose tissue.

To address if the observed increase in adipose Hif1α in HFD maintained mice contributes to the expansion of adipose tissue. Hif1α^fl/fl^ and Hif1α cKO mice were maintained on eitherNCD or HFD, and weight gain assessed over a period of 24 weeks (Fig. 2F-H). While negligible differences were detected in Hif1α^fl/fl^ and Hif1α cKO when maintained on NCD, a marked decrease in weight gain and adiposity was observed as early as 4-weeks post-HFD in Hif1α cKO compared to Hif1α^fl/fl^ mice (Fig. 2F-H). This pattern of reduced adiposity and body weight gain of Hif1α cKO mice in response to HFD continued throughout the 24-week period of nutrient overload (Fig. 2H). In spite of the reduction in WAT mass, Hif1α cKO mice did not display a lipodystrophic phenotype as evidenced by the lack of ectopic fat redistribution to peripheral tissues such as the liver or skeletal muscle in Hif1α cKO mice (Supplementary Fig. 1A, B). The liver and skeletal muscle of Hif1α cKO mice were of comparable weight and gross morphology to Hif1α^fl/fl^ mice, and did not show evidence of increased lipid content or of morphological alterations (Supplementary Fig. 1A, B).

Defective glucose homeostasis and loss of peripheral insulin sensitivity occurs as a consequence of excessive adiposity, in part due to ectopic deposition of fat in the liver and skeletal muscle. To determine the impact of both NCD and HFD on glucose homeostasis upon adipocyte Hif1α inactivation, Hif1α^fl/fl^ and Hif1α cKO mice were subjected to a glucose tolerances test (GTT). As shown in Fig. 2I, NCD maintained mice of both groups responded comparably to the GTT. In contrast, within the HFD group, Hif1α cKO mice demonstrated significantly improved glucose handling and sensitivity compared to Hif1α fl/fl mice (Fig. 2J). As differences in glucose handling can be attributed to either decreased peripheral insulin sensitivity or reduced insulin secretion, an insulin tolerance test (ITT) was performed.

Although peripheral insulin sensitivity was comparable between mice of either groups when maintained under NCD, a marked difference in peripheral insulin sensitivity was observed specifically in Hif1α cKO mice maintained on HFD (Fig. 2K, L). While Hif1α^fl/fl^ mice maintained on HFD demonstrated little response to the bolus insulin injection, blood glucose levels in Hif1α cKO mice dropped significantly in response to insulin, indicative of maintenance of normal peripheral insulin sensitivity. Taken together, these data implicate a key role for Hif1α in nutrient overload-induced adipose mass expansion and in the maintenance of peripheral insulin sensitivity.

### Adipocyte specific Hif1α ablation reduces adipocyte cell size and number in HFD induced obesity in mouse *in vivo*

We speculated that WAT Hif1α plays a predominant role in obesity development and potentially underlies the reduction in body weight detected in Hif1α cKO upon HFD (Fig. 2H). Thus, we analyzed WAT content in Hif1α^fl/fl^ and Hif1α cKO mice under HFD. As noted, visceral WAT derived from Hif1α^fl/fl^ mice under HFD were dramatically larger than that derived from Hif1α cKO mice (Fig. 3A, B). To determine whether the reduced adiposity in Hif1α cKO mice was due to fewer fat cells and/or smaller fat cells, we compared adipocyte cell size and number in Hif1α^fl/fl^ and Hif1α cKO adipocytes. As adipocytes are filled with a large intracellular pool of TAG, the cytoplasm and nucleus are restricted to the cell periphery allowing the use of phalloidin, which marks actin filaments, as a marker of the cell boundary in immunofluorescence confocal microscopic analysis of WAT sections. Hif1α cKO adipocytes were 50% smaller than Hif1α^fl/fl^ control adipocytes but were comparable in number (Fig. 3C-E). Consistent with the difference in cell size, Hif1α cKO adipocytes demonstrated reduced incorporation of the lipid marker oil red O (ORO) and TAG content compared to Hif1α^fl/fl^ adipose tissue (Fig. 3F, G).

**Figure 3.**
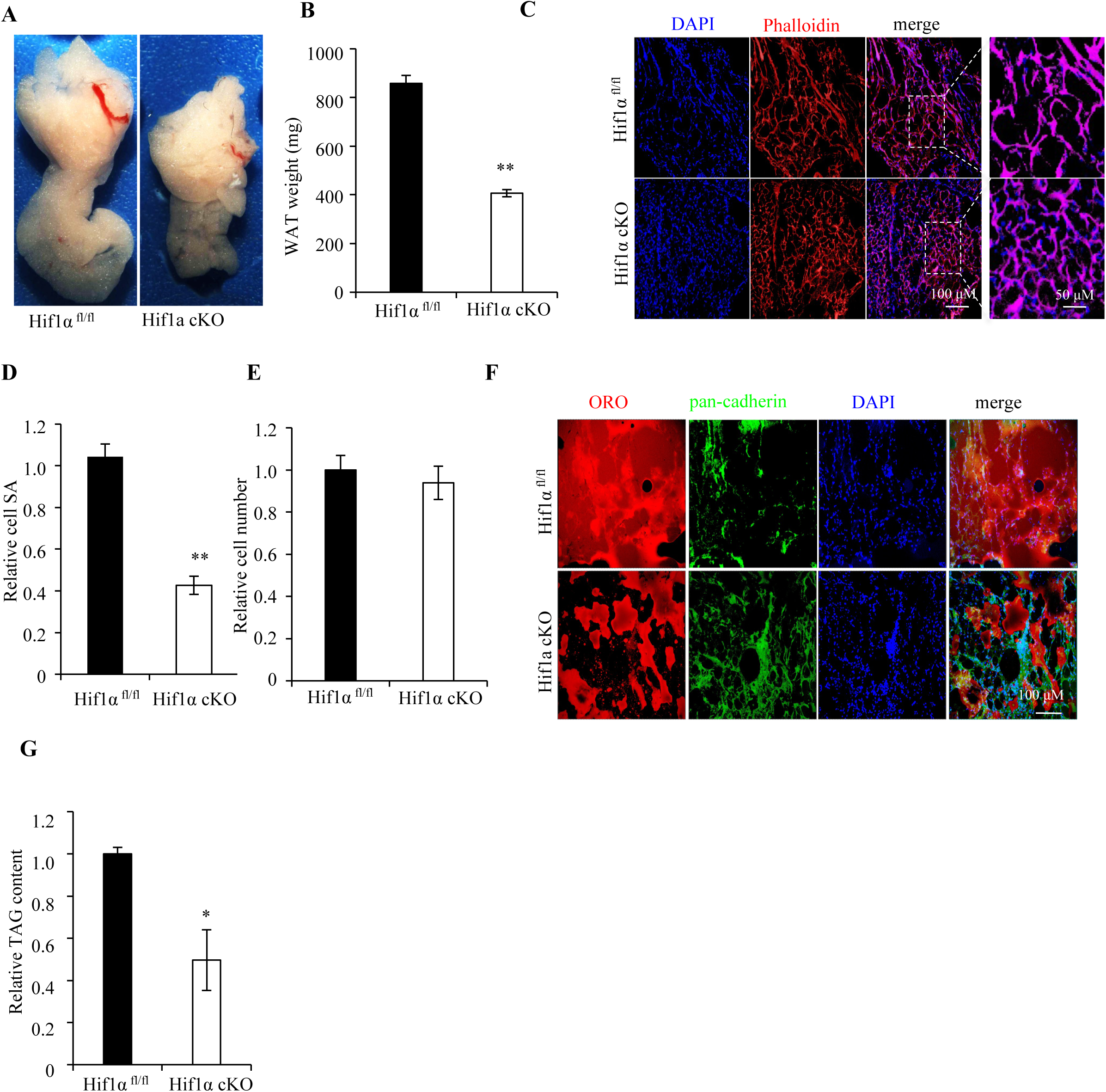
Adipocyte Hif1α inactivation reduces adipocyte cell size in HFD induced obesity in mouse *in vivo*. (A) Representative images depicting the gross morphology of WAT derived from Hif1α^fl/fl^ and Hif1α cKO mice maintained on HFD protocol. (B) Visceral WAT weight for the Hif1α^fl/fl^ and Hif1αcKO mice maintained on HFD. Data are expressed as mean ± SEM (n=6). \**P* < 0.05 vs. Hif1αfl/fl HFD, all by unpaired two-tailed Student’s t-tests. (C) WAT sections of Hif1α^fl/f^ and Hif1α cKO mice maintained on NCD were stained with DAPI (blue) and with phalloidin to mark actin (red), and analyzed by immunofluorescence microscopy. Scale Bar: 100 μM. A magnified view of the area within the dotted rectangle in the merge panel is shown on the extreme right. Scale Bar: 50 μM. (D) Quantification of visceral WAT cell surface area (SA) of HFD-fed Hif1α^fl/fl^ and Hif1α cKO mice. Data are expressed as mean ± SEM (n=6). \*\**P* < 0.01 vs. Hif1α^fl/fl^ HFD, all by unpaired two-tailed Student’s t-tests. (E) Quantification of the adipocyte cell number derived from visceral WAT of Hif1α ^fl/fl^ and Hif1α cKO mice maintained on HFD. Data are expressed as mean ± SEM (n=5). All by unpaired two-tailed Student’s t-tests. (F) WAT sections of Hif1α^fl/fl^ and Hif1α cKO mice maintained on NCD were stained with the neutral lipid marker, oil red O (ORO, red), and DAPI (blue), and counterstained with the cell boundary marker, pan-cadherin (green), and analyzed by immunofluorescence microscopy. Scale Bar: 100 μM. (G) Quantification of visceral WAT TAG content of Hif1α^fl/fl^ and Hif1α cKO mice. TAG content was normalized to tissue weight. Data are expressed as mean ± SEM (n=7). \*\**P* < 0.05 vs. Hif1α^fl/fl^ HFD, all by unpaired two-tailed Student’s t-tests. NCD, normal chow diet; HFD, high-fat diet; WAT, white adipose tissue.

### Hif1α regulates adipocyte growth through the glycerolipid biosynthetic shunt

Based on these observations we investigated if the reduced WAT growth in Hif1α cKO mice maintained on HFD occurred as a result of altered Pparγ activation and reduced flux through the glycerolipid biosynthesis pathway in adipocytes (Fig. 4A). As shown in Fig. 4B, C, RNA and protein levels of Pparγ and its gene targets Gpd1 and Gpat were reduced in Hif1α cKO WAT biopsies, compared to controls. Although the decreased levels of Gpd1 and Gpat observed in Hif1α cKO mice maintained on HFD is suggestive of decreased de novo TAG synthesis capacity, we directly assessed if flux through the glycerolipid shunt was altered in WAT of Hif1α cKO mice. To that end, WAT biopsies of Hif1α^fl/fl^ and Hif1α cKO mice maintained on HFD were assessed for Gpd1 and Gpat enzymatic activity (Fig. 4D, E). Gpd1 and Gpat are relevant markers of flux through the glycerolipid shunt by virtue of Gpd1 being the first step of the glycerolipid biosynthesis pathway and Gpat being the rate-limiting step for de novo TAG synthesis. Nutrient overload induced a significant rise in the enzymatic activity of the glycerolipid pathway enzymes Gpd1 and Gpat in Hif1α^fl/fl^ WAT (Fig. 4D, E). However, inactivation of HIF1α function in adipocytes prevented the induction of Gpd1 and Gpat enzymatic activities in response to nutrient overload (Fig. 4D, E). Thus, Hif1α regulates flux through the glycerolipid biosynthetic shunt.

**Figure 4.**
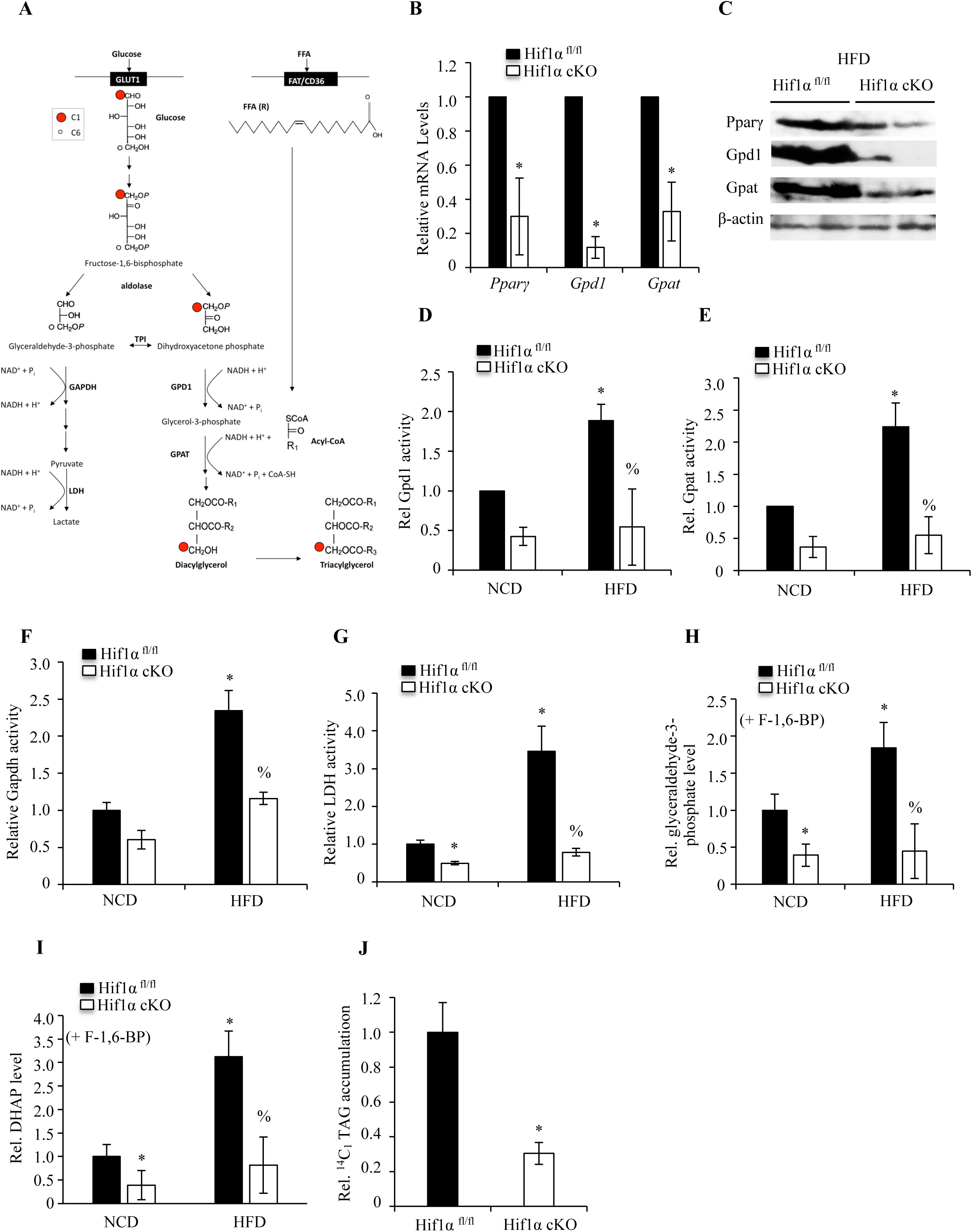
Adipocyte Hif1α inactivation reduces glycerolipid biosynthesis in HFD induced obesity. (A) Glycolytic and glycerolipid biosynthesis pathway. (B) Relative mRNA expression levels of the *Pparγ*, *Gpd1* and *Gpat* in the visceral WAT of Hif1α^fl/fl^ or Hif1α cKO mice maintained on HFD protocol. Data are expressed as mean ± SEM (n=3). \**P* < 0.05 vs. Hif1α^fl/fl^ HFD; all by unpaired two-tailed Student’s t-tests. (C) Western blots show the protein levels of Pparγ, Gpd1 and Gpat in the visceral WAT of Hif1α^fl/fl^ or Hif1α cKO mice maintained on HFD protocol. (D) Visceral WAT of Hif1α^fl/fl^ or Hif1α cKO mice were assessed for endogenous Gpd1 enzymatic activity. Data are expressed as mean ± SEM (n=4-6). \**P* < 0.05 vs. Hif1α^fl/fl^ NCD; %*P* < 0.05 vs. Hif1α^fl/fl^ HFD. Unpaired two-tailed Student’s t-tests. (E) Visceral WAT of Hif1α^fl/fl^ or Hif1α cKO mice were assessed for endogenous Gpat enzymatic activity. Data are expressed as mean ± SEM (n=4-6). \**P* < 0.05 vs. Hif1α^fl/fl^ NCD; %*P* < 0.05 vs. Hif1α^fl/fl^ HFD. Unpaired two-tailed Student’s t-tests. (F) Visceral WAT of Hif1α^fl/fl^ or Hif1α cKO mice were assessed for endogenous Gpat enzymatic activity. Data are expressed as mean ± SEM (n=3). \**P* < 0.05 vs. Hif1α^fl/fl^ NCD; %*P* < 0.05 vs. Hif1α^fl/fl^ HFD. Unpaired two-tailed Student’s t-tests. (G) Visceral WAT of Hif1α^fl/fl^ or Hif1α cKO mice were assessed for endogenous LDH enzymatic activity. Data are expressed as mean ± SEM (n=3). \**P* < 0.05 vs. Hif1α^fl/fl^ NCD; %*P* < 0.05 vs. Hif1α^fl/fl^ HFD. Unpaired two-tailed Student’s t-tests. (H) Glyceraldehyde-3-phosphate level measurements in the Hif1α^fl/fl^ or Hif1α cKO mice maintained on either a NCD or HFD protocol. Data are expressed as mean ± SEM (n=4-6). \**P* < 0.05 vs. Hif1α^fl/fl^ NCD; %*P* < 0.05 vs. Hif1α^fl/fl^ HFD. All by unpaired two-tailed Student’s t-tests. (I) DHAP level measurements in the Hif1α^fl/fl^ or Hif1α cKO mice maintained on either a NCD or HFD protocol. Data are expressed as mean ± SEM (n=4-6). \**P* < 0.05 vs. Hif1α^fl/fl^ NCD; %*P* < 0.05 vs. Hif1α^fl/fl^ HFD. All by unpaired two-tailed Student’s t-tests. (J) ^14^C1-TAG accumulation measurements in the Hif1α^fl/fl^ or Hif1α cKO mice maintained on a HFD protocol. Data are expressed as mean ± SEM (n=4-6). \**P* < 0.05 vs. Hif1α^fl/fl^ HFD. Unpaired two-tailed Student’s t-tests. NCD, normal chow diet; HFD, high-fat diet; WAT, white adipose tissue.

Glycolysis-driven fructose-1,6-bisphosphate breakdown by aldolase gives rise to dihydroxyacetone phosphate (DHAP) and glyceraldehyde-3-phosphate (G3P). Generated DHAP can either be dehydrogenated via the glycerolipid shunt to form glycerol-3-phosphate by Gpd1, or converted to glyceraldehyde-3-phosphate and returned to the glycolytic branch by triose phosphate isomerase (TPI). Accordingly, we reasoned that in the context of increased HIF1α-mediated glycolytic capacity, and increased flux into the glycerolipid biosynthetic shunt, HIF1α function could serve to ensure a coordinate engagement of the glycolytic and glycerolipid biosynthetic branches (Figure 4A), thereby simultaneously promoting the generation of glycolytic intermediates, and its channeling into and utilization by the glycerolipid biosynthetic shunt. To assess if the establishment of such a mode of co-regulation exists in adipocytes, we assayed enzymatic activity along the canonical glyceraldehyde-3-phosphate dehydrogenase (Gapdh)-lactate dehydrogenase (Ldh) branch. Gapdh activity is a key rate-limiting step in glycolysis, while lactate dehydrogenase (Ldh) serves as readout for overall glycolytic rate. As noted in Fig. 4G, H, both Gapdh and LDH activity was markedly reduced in Hif1α cKO WAT biopsies compared to controls. In accord, we also detected coordination between the canonical Gapdh-LDH axis with the Gpd1-Gpat axis in isolated primary adipocytes of Hif1α^fl/fl^ and Hif1α cKO mice, and in lipogenic NIH 3T3-L1 cells (Supplementary Fig. 2A-G). These studies not only confirm the coordination of glycolysis with glycerolipid biosynthesis specifically in adipocytes, but further reveal the plasticity of the response to Hif1α function (Supplementary Fig. 2A-G).

If HIF1α function does indeed simultaneously coordinate engagement of the glycolytic and glycerolipid biosynthetic branch (by increasing flux from glycolysis for re-channeling towards the glycerolipid biosynthetic branch), one would predict an impact of Hif1α-deficiency on relative levels of the breakdown products of the aldolase reaction (Fig. 4A). Hence, we determined if conversion of fructose-1,6-bisphosphate to either glyceraldehyde-3-phosphate or DHAP (by aldolase) was altered between Hif1α^fl/fl^ and Hif1α cKO WAT. Biopsies derived from the respective genotypes were first depleted of endogenous fructose-1,6-bisphosphate, glyceraldehyde-3-phosphate, DHAP and glycerol-3-phosphate. Thereafter, exogenous fructose-1,6-bisphosphate was added and the products of the aldolase reactions, then glyceraldehyde-3-phosphate and DHAP were measured. Nutrient overload caused enhanced production of glyceraldehyde-3-phosphate and DHAP in Hif1α^fl/fl^ WAT, while Hif1α-deletion blocked nutrient overload-induced metabolite production (Fig. 4H, I). Taken together, these data suggest a role for adipocyte Hif1α in regulating diet-induced adiposity through the co-regulation of glycolysis, and re-channeling of glycolytic intermediates into the glycerolipid biosynthetic pathway for de novo TAG synthesis.

As a result of glucose metabolism, DHAP generated from the breakdown of fructose-1-6-bisphosphate by aldolase retains the carbon moiety at position 1 of glucose (C1), which upon conversion to glycerol-3-phosphate by GPD1 is acylated by GPAT to culminate in TAG incorporation (Fig. 4A). Glucose labeled at C1 would be predicted to incorporate into TAG, to a greater extent in adipocytes of Hif1α^fl/fl^ mice maintained HFD compared to Hif1α cKO adipocytes, if indeed Hif1α promoted the preferential channeling of the glycolytic intermediate, DHAP, towards TAG synthesis and, if the glycerol component of the stored lipids did in fact derive from the glycolytic cascade. ^14^C1-glucose was added to WAT explants derived from Hif1α^fl/fl^ and Hif1α cKO mice subjected to HFD, and stored TAG extracted and analyzed for ^14^C1 incorporation. As shown in Fig. 4J, ^14^C1-incorporation into TAG was significantly reduced in Hif1α cKO biopsies, compared to controls. Thus, activation of Hif1α in response to nutrient overload contributes significantly to the conversion of glucose-derived carbon to lipid synthesis, and adipocyte hypertrophy.

## Discussion

Here we demonstrate coupling of glycolysis and lipid anabolism in the regulation of basal and diet-induced adiposity and provide evidence for the integration of the two metabolic pathways through the transcription factor Hif1α. The coupling of these pathways facilitates triacylglyceride synthesis resulting in increased lipid deposition and adipocyte hypertrophy, culminating in adipose mass expansion. Importantly, our data reveals a necessity and sufficiency for coupling of the glycolytic and glycerolipid biosynthetic pathways in both developmental and nutrient overload-induced adipose tissue growth. These changes in adiposity, resultant of Hif1α-mediated metabolic coupling, further impacts systemic metabolism through disruption of glucose homeostasis and peripheral insulin sensitivity.

Mechanistically, our work unveiled that Hif1α function leads to rewiring of adipocyte metabolism, wherein glycolytic intermediates are partially redirected into the glycerolipid shunt, thereby facilitating the de novo synthesis of triacylglycerides. Thus, Hif1α function coordinates the output of the respective metabolic pathways through simultaneous and coordinated control of both glycolysis and triacylglyceride biosynthesis. Hif1α-mediated transcriptional induction of glycolytic genes, and of Pparγ its target genes Gpd1 and Gpat, leads to increased glycolysis, and concomitant redirection of glycolytic intermediates due to activation of the glycerolipid biosynthetic shunt. Hence, coupling of these two metabolic pathways facilitates adipocyte growth and white adipose mass expansion.

We hypothesize that a secondary level of regulation with respect to NAD+ and NADH levels might serve in the maintenance of the two branches. Gapdh utilizes NAD+ in 1,3-bisphosphoglycerate generation, with NADH being a byproduct. Gpd1, in contrast, requires NADH for glycerol-3-phosphate generation with NAD+ as its byproduct. There is evidence to suggest of a physical interaction and/or coupling of NAD+/NADH moieties between the two enzyme complexes (32). Thus, continuous cycling of NAD+ and NADH between the complexes might serve to sustain a high flux through the respective branches under conditions of high intracellular glucose, as is typical of the hypertrophic, hyperglycemic, anaerobic and insulin-induced states. The latter idea is supported by our data documenting a simultaneous decrease of Gapdh and Gpd1 enzymatic activities in adipose tissue of mice maintained on HFD. Furthermore, the demonstration that the carbon moiety of exogenously added glucose is channeled, under the above-noted conditions into the lipid fraction of adipocyte underscores the critical requirement of the glycolytic and glycerolipid synthesis pathways for lipid accumulation and adipocyte hypertrophy.

It is important to note that reports of coupling between glycolysis and triacylglyceride synthesis have been noted since the 1960s. Research in the past decades reveals that low oxygen promoted glycolysis, glycerol-3-phosphate generation and lipid synthesis in adipose tissue and skeletal muscle (29,33,34). However, the underlying mechanism mediating these changes in lipid synthesis in response to hypoxic stress has remained unclear. Based on the above study, we can now propose a mechanistic explanation for the observed phenomenon in which the coordinated upregulation of glycolysis directly by HIF1α and its indirect promotion of the fatty acid uptake and the glycerolipid synthesis pathway via PPARγ brings about the metabolic shift characteristic of the diseased state.

In its totality, our work identifies Hif1α as a central regulator of white adipose tissue growth and of systemic glucose homeostasis through its capacity to integrate a central catabolic pathway with lipid anabolism. These data have implications for the use of PPARγ agonist for treatment of type 2 diabetes and suggest although beneficial for normalization of glucose homeostasis in the short tem, long-term use of PPARγ agonist may lead to deterioration of systemic glucose regulation due to increased visceral adipocyte growth.

## Acknowledgements

We thank N. Fankhauser, K. Eckhardt and S. Georgiev for scientific help and advice, and all members of the Institute for Cardiovascular Regeneration and Genome Biologics for technical, regulatory, administrative, analytical, reagent and scientific support. We are indebted to R. Johnson and R. Evans for mouse lines. This work was supported by the Medical Research Council (MRC MCUP1102/4), LOEWE Center for Cell and Gene Therapy, and the European Innovation Council (GA: 822455) to J.K., the Foundation for Pathobiochemistry and Molecular Diagnostics to P.M. Y.W. and Y.M. were supported by the China Scholarship Council (CSC) Grant #202108080020 and #202306310045, respectively.

## Author Contributions

P.M., T.Y., and J.K. planned and designed the experiments, and wrote the paper. C.Z., M.W., M.D.P., Y.W., A.P.D., Y.M. and J.K. executed most of the experiments with experimental and analytical help from T.Y., A.P.D., M.W., Y.M. and P.M.

## Supplementary Figure Legends

**Supplementary Figure 1.** Adiopocite specific Hif1α ablation does not alter the metabolic genes expression. (A) Liver of Hif1α^fl/fl^ and Hif1α cKO mice maintained on the HFD protocol were stained for the pan-cadherin (green), oil red O (ORO, red), and DAPI (blue) and analyzed by immunofluorescence confocal microscopy. Scale Bar: 200 μM. (B) Representative images of the skeletal muscle for Hif1α^fl/fl^ and Hif1α cKO mice maintained on the HFD protocol. Scale Bar: 5 μM. HFD, high-fat diet; WAT, white adipose tissue.

**Supplementary Figure 2.** Adipocyte Hif1α inactivation reduces glycerolipid biosynthesis in adipocytes *ex vivo* and *in vitro*. (A-D) Primary adipocytes were isolated from HFD-fed Hif1α^fl/fl^ and Hif1α cKO mice, and the activities of Gpd1 (A), Gpat (B), Gapdh (C) and Ldh (D) were assessed. Data are expressed as mean ± SEM (n=5). \**P* < 0.05 vs. Hif1α^fl/fl^ HFD. Unpaired two-tailed Student’s t-tests. (E-G) Lipogenic NIH 3T3-L1 cells were transduced by lentivirus carrying Hif1α shRNA or Hif1α, and then ECAR was measured with seahorse analyzer (E), Gapdh activity (F) and Ladh (G) were assessed. Data are expressed as mean ± SEM (n=4). \**P* < 0.05 vs. ns Control. Unpaired two-tailed Student’s t-tests. HFD, high-fat diet; ECAR, extracellular acidification rate.

