## Supplementary figures and images for "HIF1α controls adipose tissue growth through metabolic rechanneling"

### Supplementary Figures

**Supplementary Figure 1**

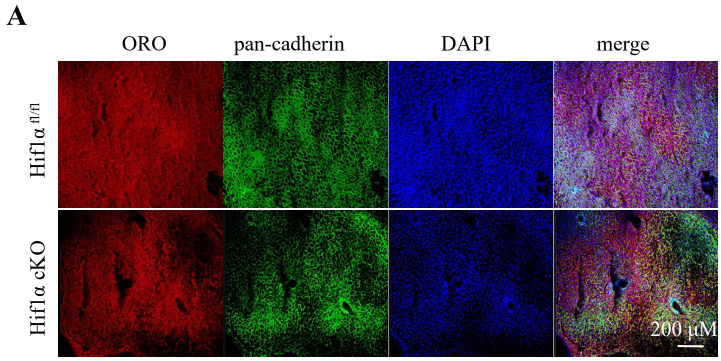


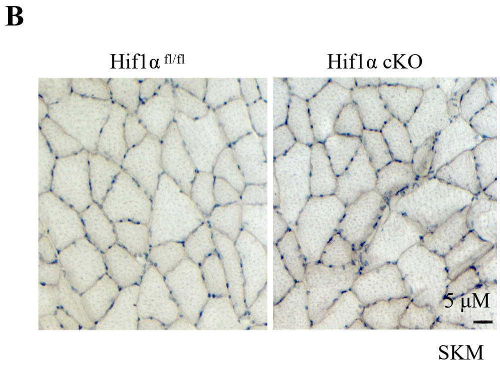

**Supplementary Figure 2**

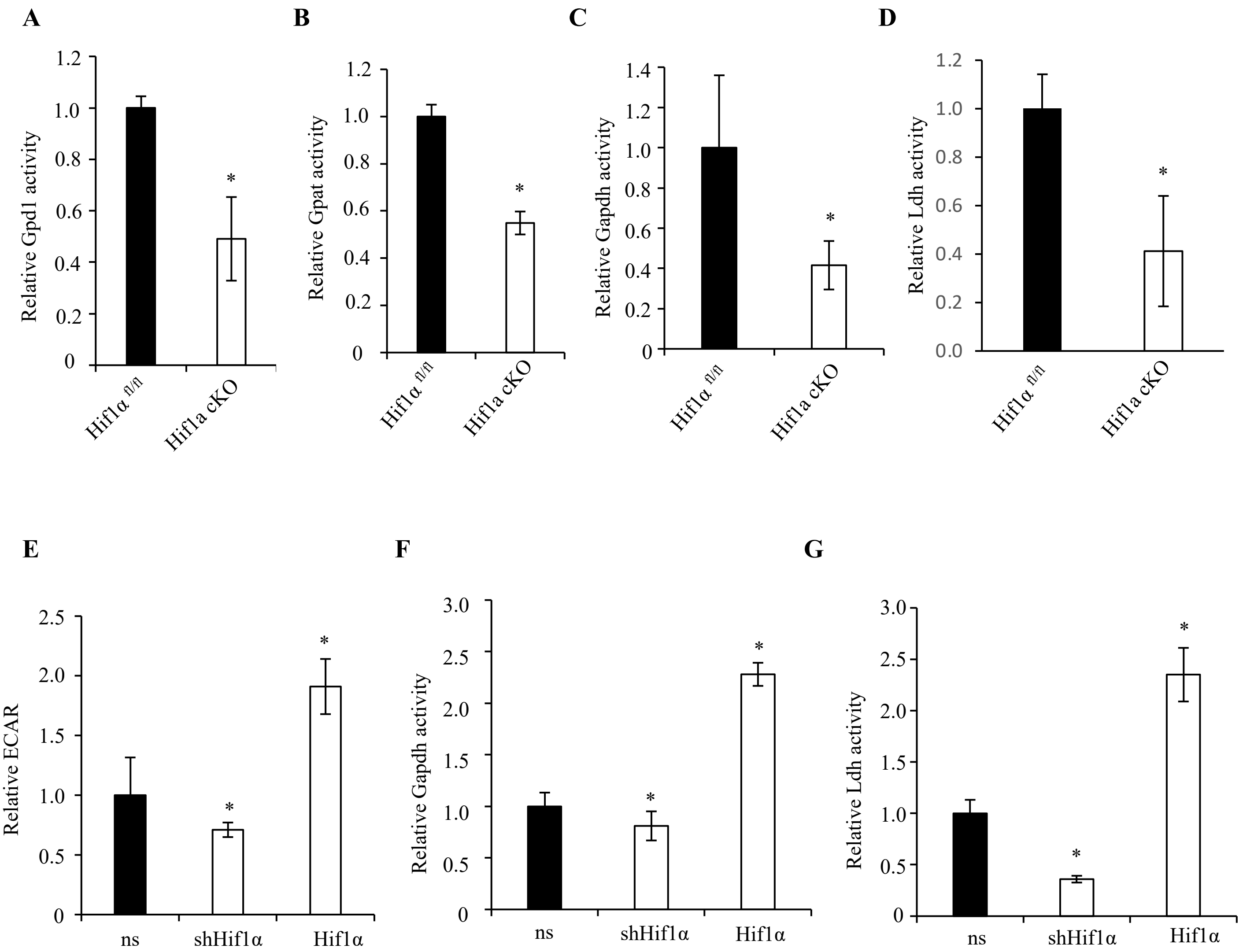
